# EEG Foundation Model Improves Online Directional Motor Imagery Brain-computer Interface Control

**DOI:** 10.64898/2026.03.24.714020

**Authors:** Maxim A. Karrenbach, Hanwen Wang, Zachary Johnson, Yidan Ding, Bin He

## Abstract

Brain-Computer interfaces (BCIs) offer a link between neural signals and external computation, enabling control of devices for the purposes of restoring function to motor-affected individuals and enhancing capabilities of a wider set of populations. Electroencephalography (EEG) offers a high temporal resolution for dynamic and potential real-time feedback for non-invasive systems. However, its practical efficacy remains limited due to low spatial resolution and poor signal-to-noise ratio, leading to insufficient decoding accuracy and unintuitive control paradigms that hinder reliable user interaction. In this study, we present a framework for an online EEG foundation model by creating a custom foundation model through spectrogram reconstruction of compact temporal windows and online constraints during pretraining. We evaluate the performance of the model in a challenging control paradigm of single-arm, directional motor imagery with dynamic movements for guided and free movement cursor control tasks. Our foundation model approach achieved a final average accuracy of 51.3% during a goal-oriented guided control task. This represents a 15.8% increase over a conventional deep learning framework and a 26.3% increase above chance level, evaluated in a cohort of 11 human participants. During the free movement task, the foundation model invoked a higher rate of completion and lower completion times. Furthermore, the custom EEG foundation model demonstrated superior adaptability from same-session finetuning and indicated an enhanced capability to assist subject learning. These findings highlight the potential of EEG foundation models to support more robust and intuitive non-invasive BCI systems, providing a promising modelling framework for future BCI development.

## INTRODUCTION

Brain-computer interfaces (BCI) are capable of providing control of external devices, with the potential to restore or provide alternative motor capabilities or communication abilities in clinical populations and beyond [1], [2]. Invasive BCI systems have created frameworks which enabled robust robotic manipulation [3], [4] or communicative speech [5], [6]. However, accessibility of such highly invasive BCIs are well limited to only severely disabled individuals due to the need for surgical procedures and brain implants, as well as the high cost. On the other hand, non-invasive electroencephalography (EEG) BCI systems have been developed to provide a safe and affordable means. These EEG-BCI frameworks have potential to benefit numerous patients to regain independence and improve quality of life, and even the general population for a variety of tasks [2].

Research studies involving noninvasive EEG-BCI have explored the capabilities of BCI to decode intent or understanding from neural signals for control of external devices and communication [7]. Early BCI studies demonstrated the ability to control a virtual cursor in 2D [8] using the sensorimotor rhythms of motor imagery (MI). Further studies extended the exploration to more complex tasks, creating frameworks to control virtual [9] and model helicopters/drones [10], as well as robotic arms [11], [12], [13], [14], [15], [16] and hands [17]. Towards clinical applications, EEG-BCI has also shown the ability to aid motor-impaired populations through control of assistive devices like wheelchairs [18], [19] or exoskeletons [20], [21]. These achievements represent the capabilities of EEG-based BCI systems for functional control in both healthy and clinical populations.

Despite that EEG suffers from a low signal-to-noise ratio, as the neural signals are attenuated between the neural activation and the scalp sensor, causing signal blurring due to volume conduction [22], progresses have been made to demonstrate the potential for noninvasive BCI control. Studies have found that some kinematic characteristics from motor execution of the upper limb [23], [24], [25], [26], [27] can be decoded, even continuously [28]. Similar characteristics have been decoded in motor imagery as well [29], [30], showing the interest in decoding naturalistic movements from EEG. However, current control methods rely on configurations of spatially distinct limbs, which can be unintuitive to subjects and limit the degrees of freedom of the system. There is a need for algorithmic advances in neural decoding for natural movement for use in BCI.

Data-driven approaches based on deep learning have emerged to offer an enhanced capacity to learn neural representations directly from minimally processed signals by capturing complex spatiotemporal patterns from EEG [31], [32], [33]. However, many existing deep learning approaches for EEG-BCI remain optimized for specific tasks and subjects, often being evaluated primarily in offline settings. Some studies have explored implementing deep learning in online settings to advance complex control [13], [17], [34], but use personalized, task-specific models and require a larger number of sessions to collect enough data to achieve high performance. Thus, the current online approaches cannot utilize general neural representations in EEG, instead optimizing via task or subject-specific characteristics which can vary due to inter-session differences in online settings. As a result, the ability of deep learning to support robust and intuitive control under the constraints of real-time BCI operation remains limited.

More recently, foundation models have been proposed as a general framework for representation learning, characterized by large-scale pretraining and adaptative fine-tuning to downstream tasks. By leveraging diverse datasets, foundation models aim to learn broad and transferable representations that extend beyond task-specific optimization. These techniques have led to substantial advances in fields such as computer vision and natural language processing [35], [36], [37]. Initial explorations which adapted foundation model techniques for offline EEG decoding [38], [39], [40] have indicated that foundation model techniques can learn generalizable neural representations from EEG from large sets of data containing a variety of tasks [41], [42], [43], and even when focusing on a subset of EEG tasks [44]. These findings suggest that foundation models may be well suited to address representational challenges inherent in EEG signals [38], [39], [40], [45].

Despite these encouraging results, the integration of EEG foundation models into online BCI systems remains largely unexplored. Real-time BCI operation imposes stringent constraints on latency, which limit the applicability of implementing existing foundation model architectures. Primarily, the detection of the subject’s action will be relayed to the system with latency dictated by the temporal length of the window used by the model. Thus, the real-time feedback of BCI mandates small temporal windows for the model, with previous studies with deep learning utilize windows ranging between one and four seconds [13], [14], [17], [34]. Additionally, online experiments often implement minimal processing or contain numerous signal changes that are not easily dealt with in real time. These constraints represent challenges which may restrict EEG foundation models from being utilized in online BCI currently.

In this study, we propose a custom EEG foundation model to address the challenges that foundation models face when incorporated into closed-loop BCI. This model uses a pretraining process which utilizes spectrogram reconstruction on small time windows to combat latency alongside minimal preprocessing. Through this exploration, we aim to assess whether an EEG foundation model can learn strong neural representations of EEG while remaining robust to the conditions of online experimentation. We evaluate this model against a conventional deep learning model in a demanding online experimental setup by using single-arm dynamic motor imagery for cursor control. Through this spatially and temporally complex paradigm, we explore the potential of online EEG foundation models to enable more intuitive and natural control in EEG-BCI through stronger neural representations guided by online BCI considerations.

## METHODS

### Experimental Design

Latency from longer time windows and a mismatch between offline data curation and online minimal preprocessing represent some of the main challenges which preclude existing EEG foundation models from incorporating into online EEG-BCI frameworks. To address these issues, we developed a *Compact Spectral-Temporal Embedding Model (C-STEM)* as a foundation model for online experimentation. Through pretraining with spectrogram reconstruction and minimal preprocessing on over 1200 hours of motor imagery human EEG data, the C-STEM framework aims to be robust in online experimentation. In addition, current online BCI control schemes can feel unintuitive and be difficult to learn and adopt, caused by an algorithmic inability to decode more naturalistic movements from EEG. We assessed the viability of the C-STEM framework through our *online foundation model experimentation* by implementing a control paradigm with single-arm dynamic directional movements for cursor control tasks, evaluating the potential that advanced foundation model techniques may hold in EEG-BCI control applications.

### Compact Spectral-Temporal Embedding Model (C-STEM)

This study introduced an architecture (Figure 1) which aims to discretize neural EEG data into embeddings from short time windows to learn underlying representations of EEG from large amounts of unlabeled EEG data. These embeddings are trained with spectrogram and raw signal reconstruction and are used in further training for task-specific downstream datasets. In pretraining, this model follows a classic encoder-decoder style architecture with a quantizer codebook for the latent space. During downstream finetuning, the encoder and codebook are frozen, and a linear layer is appended for task-specific classification.

**Figure 1.**
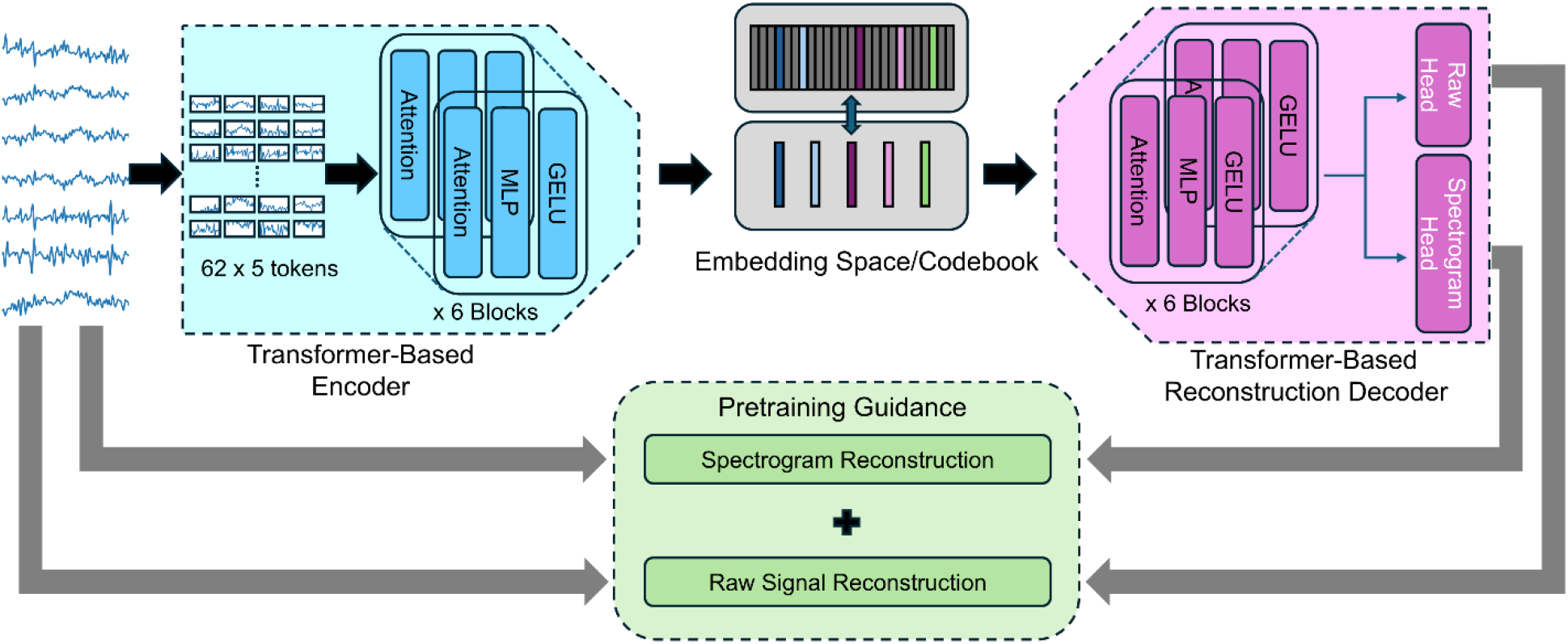
Overview of the C-STEM architecture. EEG signals are input as (time × channel) signals and flattened such that each channel is processed independently. Signals are processed in 200 ms patches. The transformed encoder and decoder act to project the signals to and from the embedding space, while the space itself is represented by a codebook of embeddings which are updated during training. Training is guided by loss terms comparing the ground truth spectrogram and raw signal to the reconstructed terms.

The key components of the architecture lie in a transformer-style [46] encoder which encodes EEG channels into the embedding space, a quantizer-codebook pair [47] to quantize, store, and replace EEG embeddings, and a decoder which mirrors the encoder in architecture and decodes raw and spectral representations of the input EEG channels from the transformed embeddings.

#### Model Architecture and Configuration

Due to the high temporal resolution of EEG data, information in both the temporal and spectral domain is key to creating discriminable features in downstream EEG tasks. Frequency domain characteristics are often used to identify motor intent via Event-Related Desynchronization/ Synchronization (ERD/ERS) [48] over the entirety of a trial. Motivated by the goal of identifying dynamic movements and the use of sliding window techniques in downstream online tasks, the architecture turns to spectrogram representations. The decoder thus attempts to reconstruct the spectrogram and raw signals from the embeddings to train the architecture to learn relevant temporal and spectral characteristics for the embeddings. The raw reconstruction loss is calculated with mean-squared error (MSE), while the spectrogram uses a weighted MSE, assigning stronger weights to the alpha (8-13Hz) and beta (13-30Hz) bands of the spectrogram for use in sensorimotor rhythm BCIs. These weights were determined to conform to the downstream tasks using motor imagery, which shows strong signatures in these bands. The total loss guiding the training is a weighted sum of the two MSE terms.

#### Pretraining and Finetuning

The model configuration details were created to closely match those details found in online experimentation. Thus, a patch size of 200ms was set for the architecture to process the 1000Hz EEG data. During pretraining, the number of patches was kept at 310, representing 1 second of EEG data split into 5 segments over all 62 available channels. When preprocessing data for pretraining, we performed only those preprocessing steps which are employed during the online experimentation. This consists of filtering data from between 0.5 Hz and 100 Hz to keep all relevant bands in the data and performing z-score normalization. Calculated spectrograms were also normalized to avoid exploding gradients.

For online experiments and any offline analysis, finetuning of C-STEM was performed by freezing the weights of the encoder and codebook, then transferring the frozen weights to an architecture fitted with a single linear layer for classification. The data is filtered identically to the pretraining process and z-score normalized. The data is also reduced down to the 27 electrodes surrounding the motor cortex (CP, C, FC according to the modified international 10-20 system). Self-supervised learning was then performed to further train C-STEM on the specific classification task relevant to the designed experiments. During online experimentation, this model was frozen and used for inference.

Pretraining and finetuning were performed on eight NVIDIA Tesla V100 GPUs using Python 3.9.19 and PyTorch 2.5.0 with CUDA 12.4 support, while inference was carried out on a NVIDIA GTX 1080-TI using Python 3.8.4 and PyTorch 2.4.0 with CUDA 11.8 support.

#### Datasets for Pretraining

The pretraining datasets (Table 1) used were open-source EEG datasets, primarily consisting of upper-limb motor imagery experimentation, with a total of 146 human participants. These datasets provide diversity in the motor imagery (MI) control paradigms, containing left/right hand MI [34], [49], finger MI [17], and foot MI [14], as well as task diversity, such as cursor control [14], [49], continuous movements [34], robotic control [14], [17] and hybrid visual attention paradigms [50]. For each of the datasets, individual trials were not extracted, rather the entire session of data recorded was utilized for training. This approach of including motor imagery signatures, along with the rest, preparation, and cue states was designed to allow for the architecture to be able to recognize the various underlying EEG representations from not only the modality of interest, but also for experimentation information. Training and test sets were created by selecting 20% of subjects from each dataset to use for testing and the remaining subjects for training. Hyperparameters for pretraining and finetuning were empirically identified.

**Table 1.** Descriptions of datasets used in the pretraining of C-STEM. Datasets used contain motor imagery experimentation and labels are available, though not used during the pretraining process.

| Dataset | Duration Used | Subject # | Tasks Performed |
| --- | --- | --- | --- |
| Stieger et al [49] | 596.8 Hours | 62 | Two-hand MI, 2D cursor control |
| Forenzo et al [34] | 162.4 Hours | 28 | Two-hand MI, Continuous 2D cursor pursuit |
| Forenzo et al [14] | 53.7 Hours | 10 | Two-hand MI, Foot MI, 2D cursor control |
| Ding et al [17] | 274.0 Hours | 21 | Single-arm Finger-level MI, robotic control |
| Forenzo et al [50] | 108.2 Hours | 25 | Two-hand MI, Overt spatial attention, 2D cursor and robotic control |

### Online Foundation Model Experimentation

We designed a complex control paradigm which utilizes motor imagery of the right arm performing single-shot directional movements and evaluated the performance of C-STEM for online BCI control in a cohort of 11 human participants. The BCI tasks focus on cursor control, wherein subjects are tasked to imagine directional movements in one of the four cardinal directions to move the cursor accordingly. Visual feedback was given either through explicit color indications of model predictions or through actual cursor movements. Two models are used randomly and blindly: a conventional deep learning model (EEGNet) and our proposed foundation model (C-STEM).

#### Single-Arm Directional Movement Offline Paradigm (Session 1,2)

Subjects first performed a paradigm where direction cues were given (Figure 2C) following a path. Subjects were instructed to concentrate on a circular cursor on a gridded screen, and a square in one of the cardinal directions around the cursor would change color. The subjects then performed an imagined or actual directional movement (Figure 2A) that would move the cursor to the indicated square. Subjects repeated this process until their circle stimulus reached a specific goal square marked with a pattern. Each path from the start to the goal was 10 spaces, and paths were verified to contain an even distribution of movements. Each movement trial lasted 3.5 seconds. The cue stage lasted 1 second, while subjects were given 1.5s to perform motor imagery (MI – imagined directional movement) or motor execution (ME – actual directional movement), with the wait stage being 1 second. Subjects were instructed to perform self-paced movements and wait for the reset stage to return to a neutral position. Each run consisted of 8 paths; Session 1 had 4 runs of MI and 1 run of ME, while Session 2 conducted 5 runs of MI and 2 runs of ME.

**Figure 2.**
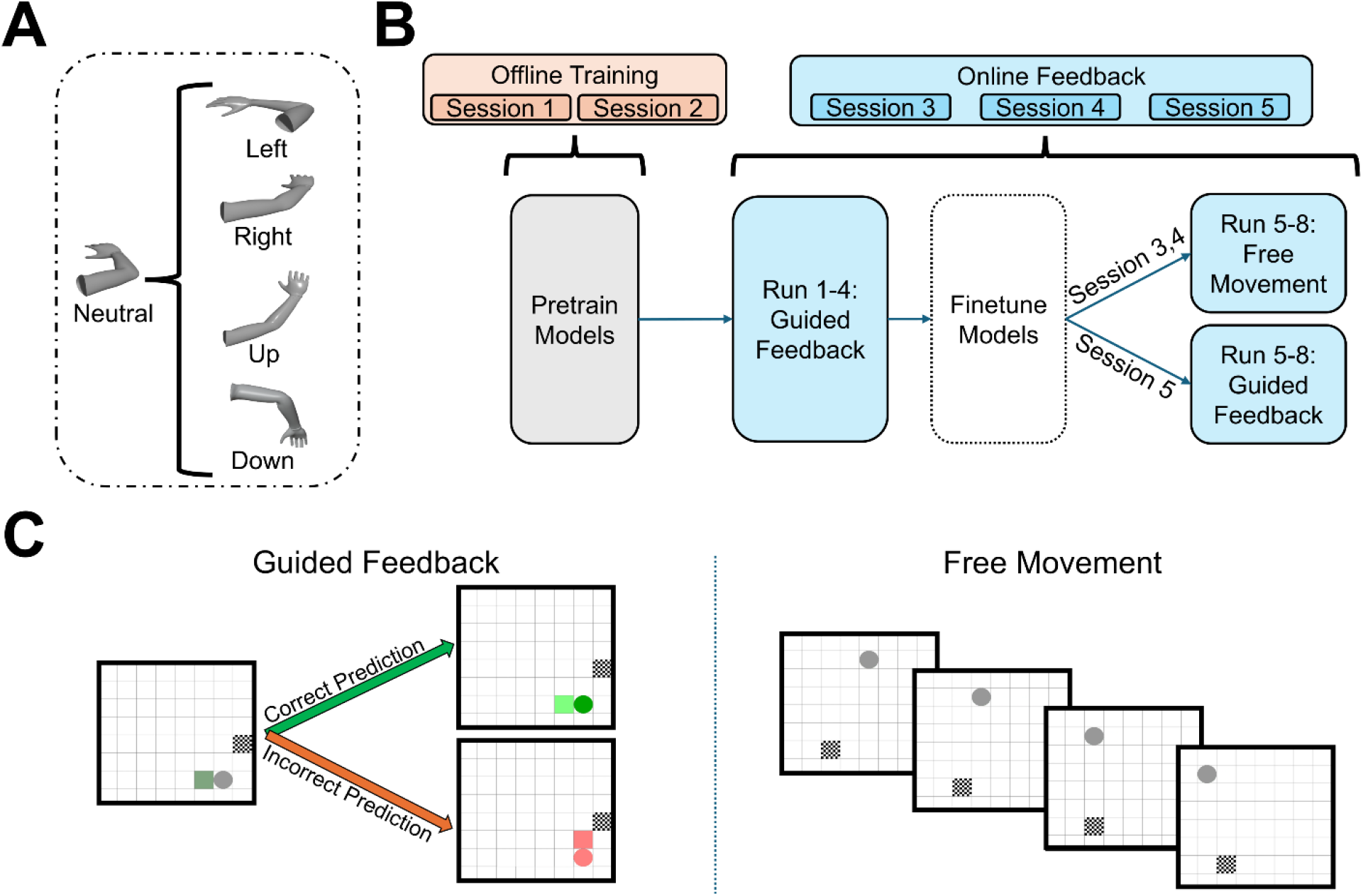
Overview of the experimental paradigms. (A) Directional movement paradigms, with subjects performing directional movements from a neutral position. (B) Experimental schedule; with the first two sessions as offline sessions, and the last three sessions containing online experimentation. Additionally, in the first half of all online sessions, subjects performed the guided movement task with a model trained on previous session’s data. In the second half, subjects run either free or guided movement sessions with a finetuned model trained on same-session data. (C) Visual descriptions of the guided movement task (left) and the free movement task (right). The guided movement task gave color-based feedback from model predictions, while the free movement task moved the cursor in the predicted direction.

#### Single-Arm Directional Motor Imagery Online Paradigm (Session 3,4,5)

In the third and fourth sessions of the experiment, subjects followed two main paradigms designed to provide online feedback.

The first paradigm was similar to the paradigm from earlier sessions, with an additional feedback phase, and is referred to as the guided movement task. The cue stage is reduced to 0.5 seconds, while the movement phase remains at 1.5 seconds with the added feedback phase for 0.5 seconds, and the wait phase kept at 1 second. These changes were made to keep total movement times at 3.5 seconds. During the subject’s movement phase, the model outputs were processed to produce a single prediction, after which the feedback phase would highlight the predicted square. If the prediction aligned with the cued direction, the square would turn green; otherwise, the square would turn red. This provided visual feedback to the subject, who could then modulate their responses if necessary.

The second paradigm developed was visually similar but removed all cues and moved the cursor according to the model’s prediction of the subjects’ motor intent and is referred to as the free movement task. Each trial is comprised of a movement phase, during which the subject performs the imagined movement, and a prediction phase, during which the processed outputs of the model are processed to a single prediction and the circle stimulus is moved accordingly. The movement phase is 1.5 seconds, followed by a wait state of 1.0 seconds. All starting and goal squares were exactly 10 moves away, and due to the possibility of subjects never reaching the goal due to difficulty, a maximum of 22 attempts was given per path.

Models used during the online sessions are trained on all data collected from previous sessions. During Sessions 3 and 4, the guided movement task was run first with four runs per session, followed by a break. During the break, the models fine-tuned with the data collected in these first four runs. Following the break, the free movement task was run with the updated model, again with four runs per session. In Session 5, four runs of the guided movement task are performed before a break. During the break, the models are finetuned with the respective labelled data from these four runs. Then, another four runs of the guided movement task are run using the updated model.

All sessions used two types of models, a conventional deep learning model, EEGNet [31], and a model of our proposed C-STEM foundation model. Before the break in each online session, each model is used for two runs each of model testing. Data collected during these test runs are labelled, so the data is processed trial-wise and used to finetune and update the models. After the break, the updated models are used for model testing in another two runs each. The choice of model during the runs is randomized and equally distributed across the models, for an equal comparison across the two models. The choice of model was not presented to the subject to avoid any biasing behavior.

#### Online Decoding and Prediction Processing

Online signal processing and model inference were performed using Python 3.8.4 scripts in the BCPy2000 library, allowing the EEG data to be passed to the model and outputs to be processed to give visual feedback or control. Every 40ms, the latest 1 second window of EEG data was processed. The 64-channel EEG signal was reduced to a 27-channel signal, by preserving only the electrodes surrounding the motor cortex (CP, C, FC according to the modified international 10-20 system). The signal was filtered between 0.5Hz and 100Hz using a fourth-order Butterworth filter, allowing a broadband signal to be preserved. The resulting temporally and spatially filtered data window is then z-score normalized and used as the input to the online inference models.

The outputs of inference are recorded for the duration of the movement phase of the subject. At the end of the movement phase, there are 37 predictions, corresponding to 1.5 seconds at an interval of 40ms. To allow enough processing time during the short trial times, the system begins processing the window predictions after approximately one second of predictions, creating a single prediction output. This post-processing finds the longest unbroken string of one class prediction and assigns this class as the predicted class for this trial. Due to the dynamic and self-paced nature of the movements, the calculation of a single prediction aimed to focus on finding the presence of a dynamic signature immediately after motor onset among the rest of the signal in the trial.

The two models used for online decoding were trained in similar manners, with each model being pretrained on all previous session data for 100 epochs (EEGNet) or 50 epochs (FM). Early stopping was employed during the training process, with a learning rate of 0.0001 (EEGNet) or 0.00001 (FM) and a weight decay of 0.001. For the EEGNet models, this finetuning process continued training on all layers and parameters, while the FM finetuning continued training most of the layers except for the quantized codebook.

#### Participants

11 able-bodied participants (mean age: 23.9±2.12, 6 male/5 female) were recruited from the greater Pittsburgh area. All experimental protocols were approved by the Institutional Review Board of Carnegie Mellon University (IRB protocol number: STUDY2017_00000548). Participants provided written informed consent. For any video recorded, participants filled out a voluntary additional video release form. All participants were considered non-naïve to EEG-BCI, since they had completed some EEG-BCI experiments previously, though the extent and the scope of the experiments were not uniform across all participants. Participants completed five sessions in total, with the last three sessions comprising online paradigms.

### Statistical Analysis

Statistical testing was conducted using Python 3.9.19 scripts and the *scipy*.*stats* and *statsmodels* libraries. To identify group differences and to measure performance comparisons between models, two-way repeated-measures ANOVA tests were run, followed by post-hoc Wilcoxon-signed rank tests with Holm-Bonferroni corrections if significant effects were found. These comparisons were used for all performance metrics in the guided movement task, including the accuracy, precision and recall metrics, as well as the created metrics of number of hits and completion time in the free movement task. To quantify specific effect sizes, Cohen’s *d* values are calculated for a subset of metrics.

## RESULTS

This study was designed to investigate the capabilities of an online-constrained foundation model to enable more complex control through improved decoding during EEG-BCI experiments. We first analyzed the decoding performance against a conventional deep learning algorithm to examine the ability of foundation models to enable more robust control through decoding performance improvements in a challenging online experiment. To demonstrate how C-STEM addresses challenges facing current foundation models, we performed analyses focusing on the model performance under latency constraints in online and offline settings. We also analyzed C-STEM’s impact on subject adaptation to the control paradigm during online experiments. Finally, we examined characteristics of C-STEM’s neural representations and the generalizability towards out-of-distribution tasks.

### Online Foundation Models (C-STEM) create higher BCI performance across tasks

We investigated the impact of using our proposed EEG foundation model in an online setting by analyzing the directional motor imagery performance during guided and free-movement tasks. C-STEM showed a significant improvement in all metrics over EEGNet during the guided movement task (Figure 3A) and the free movement task (Figure 3B). In the guided movement task, C-STEM achieved a final accuracy of 51.3% in the final session (Supplementary Figure 1) and achieved a mean accuracy over all online sessions of 47.5%. Alternatively, EEGNet reached a peak accuracy of 35.5% in the final session and a mean accuracy of 33.0% over all online sessions. During the free movement task, C-STEM achieved an average of 3.97 hits with an average completion time of 33.8s, while the EEGNet model reached a lower metric performance, reaching an average of 2.75 hits and an average completion time of 37.4s.

**Figure 3.**
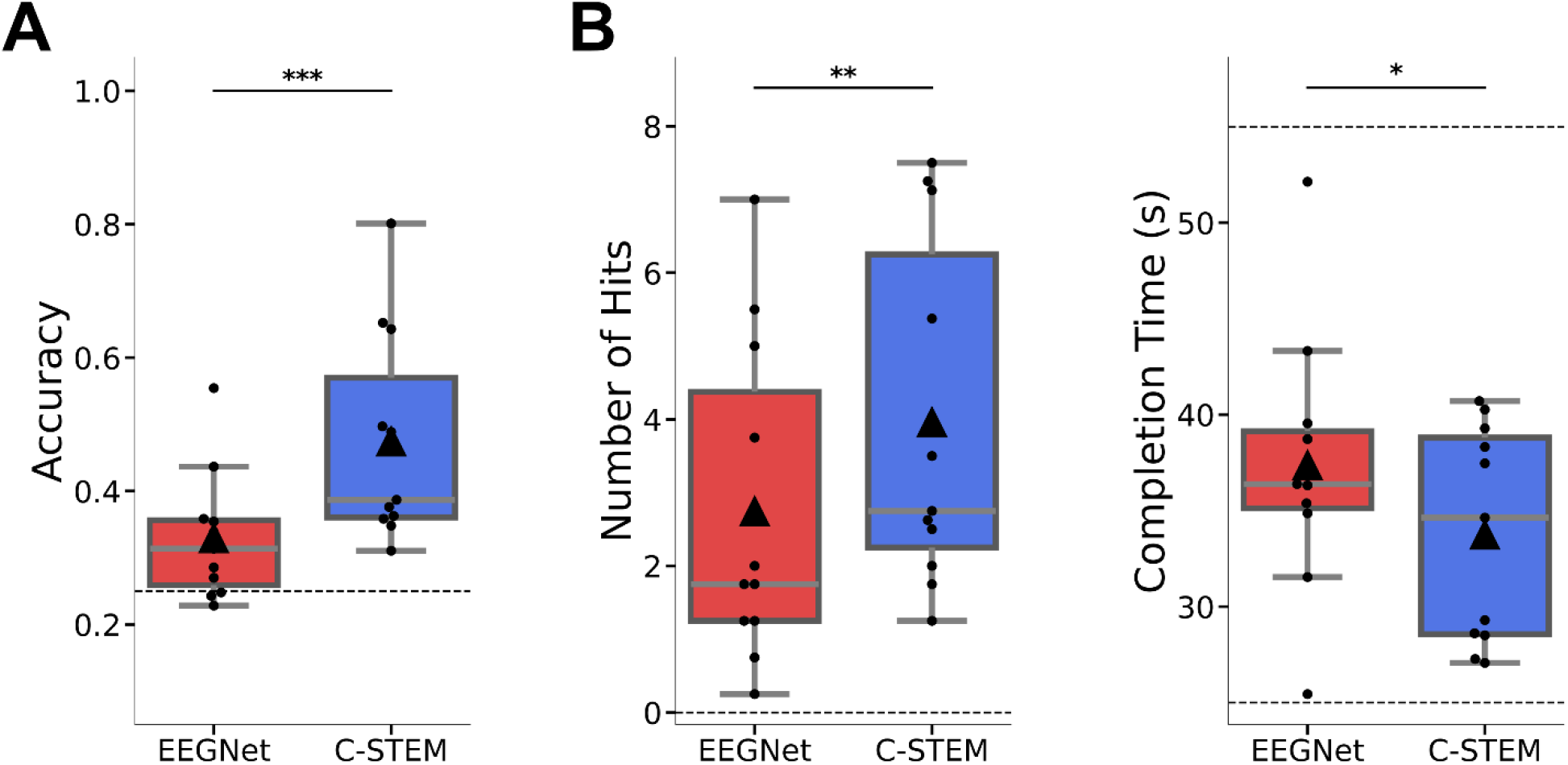
High level performance metrics averaged over all online sessions. (A) Box-plot accuracy comparisons across models displaying average accuracy over the three online sessions for the guided movement task. Center lines indicate median values, triangle markers indicate mean values, and boxes extend from the lower to upper quartile. Significance testing results between groups are displayed above bars (*** for p<0.001, ** for p<0.01, * for p<0.05, n.s. if no significance found). (B) Box-plot comparisons across models displaying the average number of hits and the completion time metrics over the three online sessions for the free movement task.

These improvements in performance metric were validated by running two-way ANOVA tests, which indicated a significant interaction between the metric and decoder architecture. These tests were conducted for each metric and revealed consistent interactions. Post-hoc Wilcoxon signed-rank tests comparing the models showed that the average accuracy (p = 0.0057), number of hits (p = 0.0088) and completion time (p = 0.024) metrics were all significantly improved under the C-STEM framework. There was a large effect size of model choice observed for accuracy (Cohen’s d = 1.099), and moderate effect sizes of model choice observed for number of hits (Cohen’s d = 0.528) and completion times (Cohen’s d = -0.586).

Furthermore, to examine the underlying behaviors responsible for the improvements, we calculated additional metrics of precision and recall and identified class-specific differences in the model performances (Figure 4, Supplementary Figure 2, Supplementary Figure 3). The improvements from C-STEM span both precision and recall averaged across classes, and confusion matrix analysis reveals that the most significant changes originate from disentanglement of the most frequently confused classes. In particular, C-STEM achieved an average precision of 0.47 and recall of 0.50, while EEGNet averaged a precision of 0.28, and a recall of 0.33.

**Figure 4.**
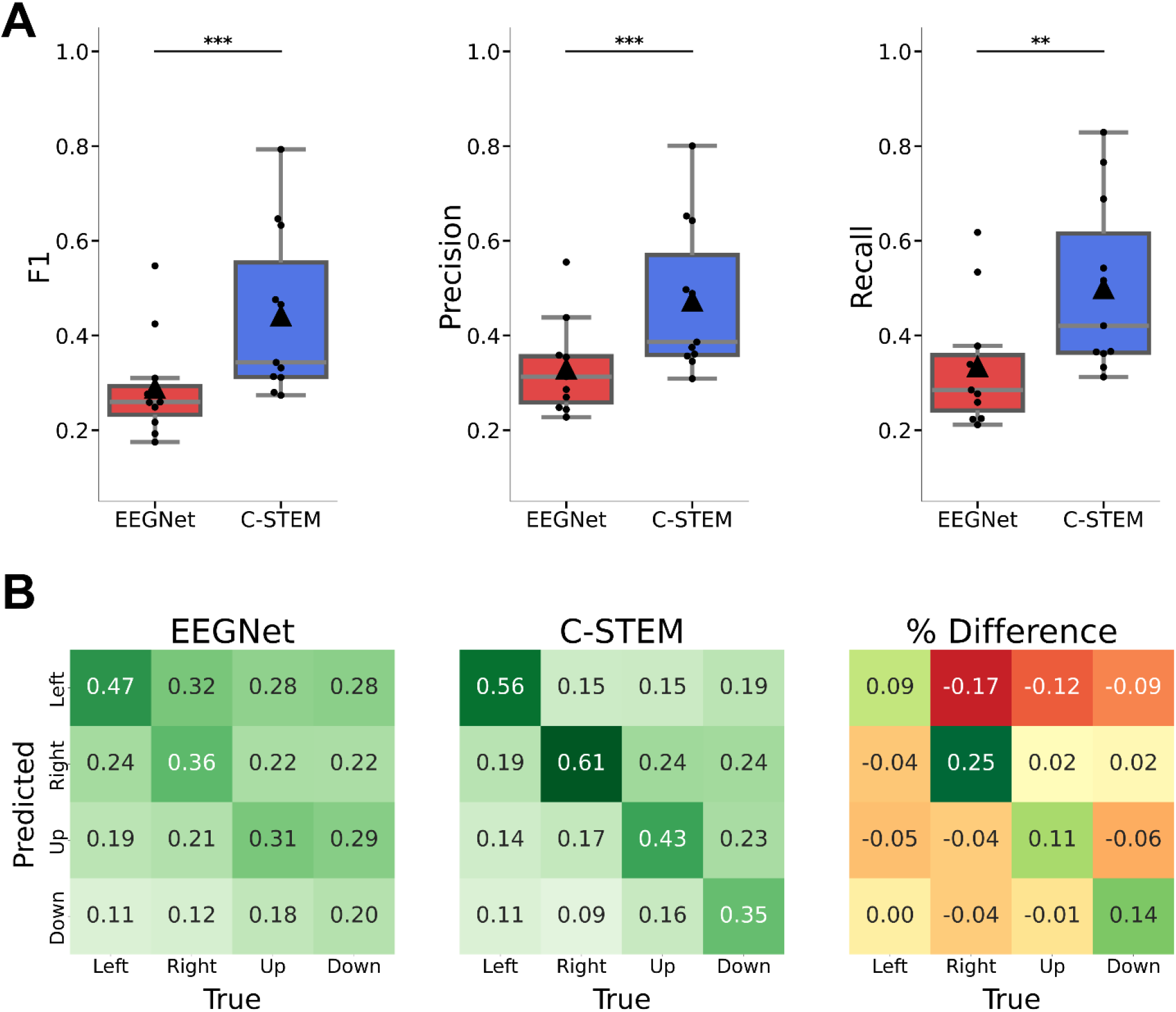
Additional performance metrics for the guided movement tasks of the online sessions. (A) Box-plot comparisons of F1, Recall and Precision metrics across models displaying average values over the three online sessions for the guided movement task. Center lines indicate median values, triangle markers indicate mean values, and boxes extend from the lower to upper quartile. Significance testing results between groups are displayed above bars (*** for p<0.001, ** for p<0.01, * for p<0.05, n.s. if no significance found). (B) Confusion matrices calculated across all subjects separated by model configuration (EEGNet on the left, FM in the center), with the percent difference between models displayed on the right. Values were normalized against the truth values and share the same scale for visualization.

Two-way ANOVA tests indicated significant effects of decoding architecture for these metrics, and post-hoc comparisons corroborated that the C-STEM metric values were significantly greater than EEGNet in both precision (p = 0.0007, Cohen’s d = 1.088) and recall (p = 0.0009, Cohen’s d = 1.029). Much of the improvements were generated from an improved ability to predict the right (+25% accuracy, p = 0.0043) and down (+14% accuracy, p = 0.048) classes.

These online results jointly demonstrate that the stringent pretraining constraints created a stronger decoding performance, stemming from an ability to better disentangle complex single-arm movements, translating to more precise control in the online setting. Together, these findings highlight the improved performance enabled by the foundation model framework in a demanding control paradigm across complex tasks.

### Online Foundation Models demonstrate lower latency in online BCI

To evaluate the impact of foundation models on decoding latency, offline analyses were performed using data collected during the online sessions. Sliding-window inference during the motor imagery phase revealed that C-STEM reached peak classification accuracy earlier than EEGNet (Supplementary Figure 4A). Specifically, peak accuracy for C-STEM occurred at 560 ms, about halfway through the trial, and achieved about 90% of the peak performance as early as 160 ms. Alternatively, EEGNet achieved peak performance at approximately 1100 ms, near the end of the trial, with the 90% peak performance reached at 840 ms. This indicates that the C-STEM framework is able to extract discriminative information from smaller temporal segments of EEG data.

To further assess performance with limited temporal information, we evaluated decoding accuracy using shorter fixed windows (200 ms, 800 ms, and 1000 ms) for multiple foundation architectures, including LaBraM [38] and NeuroGPT [51], alongside our proposed model (Supplementary Figure 4B). While all models exhibited reduced performance with 200 ms windows, our C-STEM outperformed both LaBraM and NeuroGPT on the smaller 200ms windows using our complex paradigm. This performance advantage was preserved for intermediate (800 ms) and longer (1000 ms) windows.

These findings indicate that incorporating short temporal windowing constraints during pretraining enhances the model’s ability to operate effectively under low-latency conditions, supporting the suitability for real-time BCI deployment.

### Online Foundation Models facilitate strong subject learning during BCI control tasks

To investigate the ability of subjects to adaptively generate discriminable data, we first performed a mixed model-data analysis (Supplementary Figure 5) to disentangle the impact of the model’s decoding performance from the model’s influence on subject adaptation. We define data subsets by the model used during the run (EEGNet-data and C-STEM-data) and use the frozen C-STEM and EEGNet for inference. The C-STEM on C-STEM-data (47.5%) significantly outperformed (p = 0.0058) the C-STEM on EEGNet-data (39.3%), while the EEGNet model on C-STEM-data (35.9%) improved (p = 0.029) over the EEGNet model on EEGNet-data (33.0%). A two-way ANOVA revealed a significant effect of mixed model-data configuration. Subsequent post-hoc comparisons revealed that the only groups not significantly different were the EEGNet on C-STEM-data and C-STEM on EEGNet-data (p = 0.64).

Next, to assess any online adaptation effect from the same-session distributions generated by the subject, we evaluated the impact of same-session finetuning during the guided movement task in Session 5. C-STEM demonstrated a significant improvement in online decoding accuracy after finetuning, whereas EEGNet showed no meaningful gain (Figure 5). Using C-STEM, mean accuracy increased from 47.4% before finetuning to 55.1% after finetuning (p = 0.023), corresponding to an average improvement of 7.90%. In contrast, EEGNet accuracy remained unchanged (34.9% before vs. 35.1% after; p = 0.89), with a negligible mean improvement of 0.13%.

**Figure 5.**
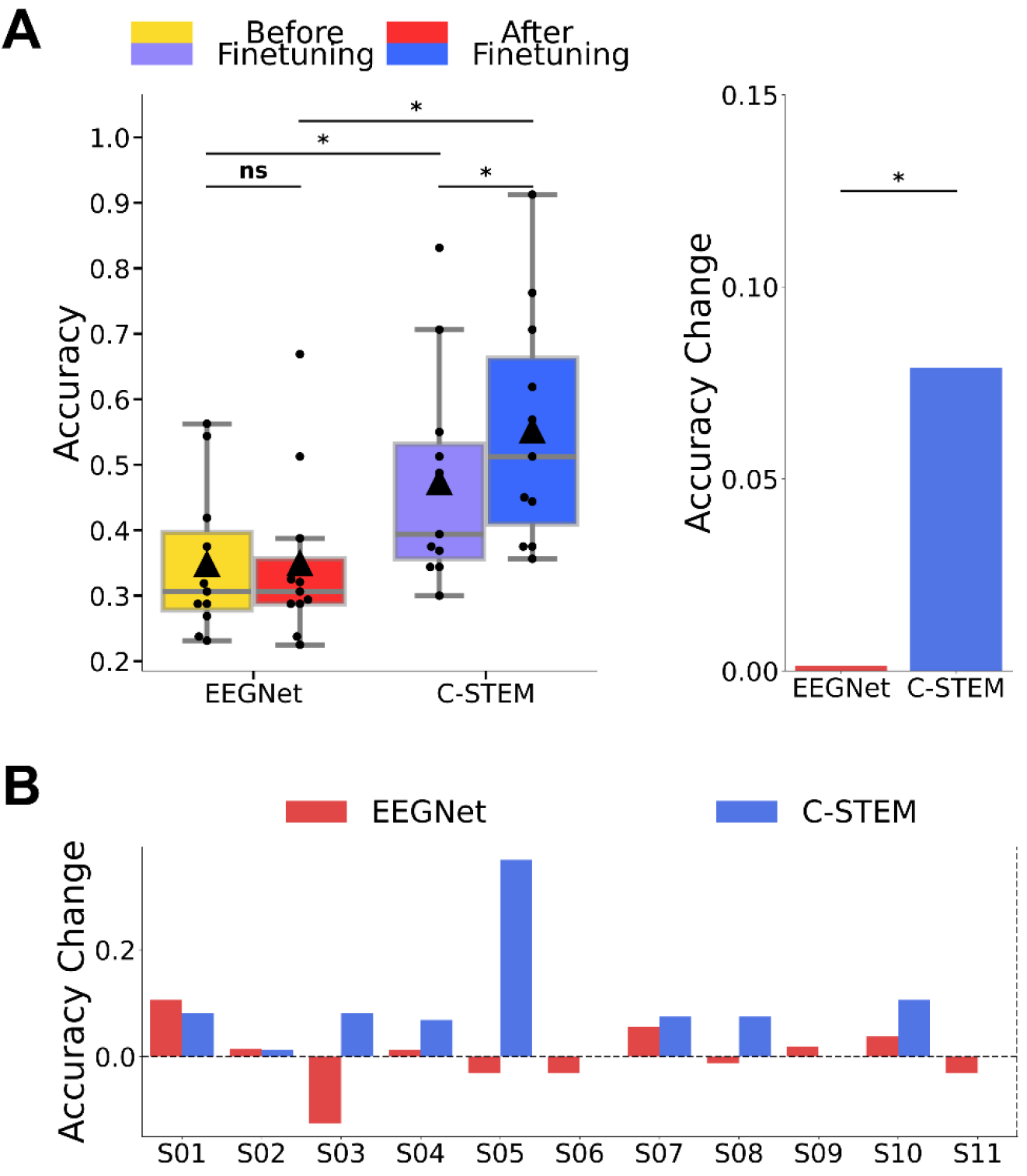
The effect of finetuning halfway through Session 5, shown on each model. (A) Box-plot visualizations of the improvements gained by finetuning for each model. Center lines indicate median values, triangle markers indicate mean values, and boxes extend from the lower to upper quartile. Significance testing results between groups are displayed above bars (*** for p<0.001, ** for p<0.01, * for p<0.05, n.s. if no significance found). The average accuracy change is shown to the right. (B) Bar plot visualizations of the accuracy change at an individual subject level.

A two-way ANOVA revealed a significant interaction between decoder architecture and finetuning condition, indicating that the performance gains were architecture dependent. Direct comparison of improvement magnitudes confirmed that the increase achieved by C-STEM was significantly greater than that of EEGNet (p = 0.024). The effect size for finetuning with C-STEM was moderate (Cohen’s d = 0.453).

These results collectively indicate a bidirectional benefit of the foundation model framework, where the subjects generate distinct session-specific data distributions under online conditions, to which C-STEM can adapt to more effectively.

### Custom C-STEM architecture creates generalizable neural representations

As the proposed C-STEM architecture focuses on short EEG temporal windows to generate embeddings through an encoder-decoder transformer block, its primary contribution lies in the neural representations it produces. To evaluate the quality of the representations learned from the architecture and shaped by the pretraining constraints in C-STEM, we first visualize the pretraining metrics. The training loss curves (Figure 6A) indicate that the pretraining conditions allow for sufficient learning of both temporal and spectral information. We also visualize reconstructions (Figure 6B) and find that these embedding representations preserve much of the low-frequency content in the signals. To visualize the utility of the embeddings created, we also compute the T-distributed Stochastic Neighbor Embedding (tSNE) [52], a non-deterministic, non-linear dimensionality reduction method for visualizing high-dimensional datasets, for both the C-STEM and the EEGNet model for one of the downstream finetuning tasks (Supplementary Figure 6). These visualizations demonstrate that the embeddings generated from C-STEM may be more separable than those of the conventional EEGNet model, leading to increased downstream performance.

**Figure 6.**
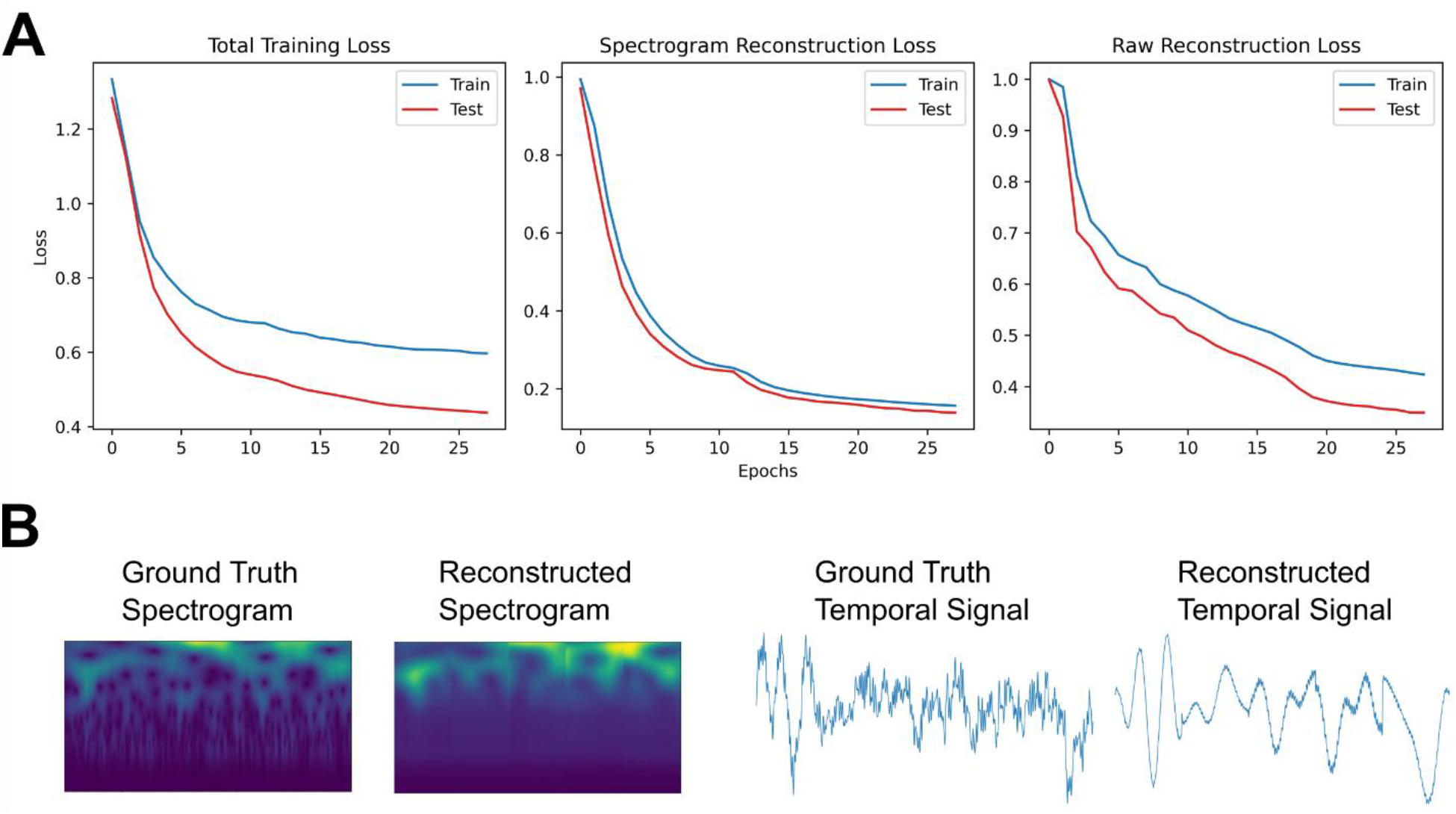
Visualizations of qualitative (reconstructions) and quantitative (loss curves) aspects of the C-STEM pretraining phase. (A) Display of loss components during pretraining over epochs. The leftmost curve displays the total training loss, as a weighted combination of the spectrogram reconstruction loss (center) and the raw reconstruction loss (right). Both training and testing curves are shown for all components. (B) Visualizations of the spectrogram reconstruction and the raw reconstruction from the proposed C-STEM pretraining process. Raw EEG signals are fed into the C-STEM architecture and outputs are calculated from the two heads of the decoder. On the left, a sample ground truth and reconstructed spectrogram are displayed, while the right shows the temporal EEG signal of this sample and the reconstructed temporal signal.

To verify the utility of the representations for downstream tasks outside the pretraining distribution, we analyzed offline motor imagery and motor execution data collected during Sessions 1 and 2. C-STEM showed significant improvements over EEGNet in both motor imagery (Supplementary Figure 7A) and motor execution (Supplementary Figure 7B) averaged across all subjects. C-STEM achieved an average accuracy of 57.6% in motor imagery and 53.8% in motor execution, while EEGNet achieved 52.7% and 49.0%, respectively. A two-way ANOVA revealed a significant effect from the decoder architecture, and post-hoc accuracy comparisons revealed that the improvement in accuracy was significant in both motor imagery (p = 0.024) and motor execution (p = 0.0068).

These results indicate that the proposed C-STEM architecture and pretraining constraints produce information-rich embeddings that enhance separability in downstream tasks, yielding neural representations that generalize across tasks and modalities.

## DISCUSSION

This study presents a foundation model framework (online C-STEM) that advances motor imagery brain-computer interfaces by designing online-constrained foundation models for decoding in the BCI loop. This approach provides insight into the potential of foundation models to increase complexity of control and tasks in BCI systems. By incorporating online constraints to guide the pretraining of C-STEM, we establish a framework that translates the demonstrated offline advantages of foundation model principles [40], [51], [53], [54] towards online BCI applications. After three sessions, subjects achieved a final accuracy of 51.3% in a two-dimensional cursor control task using a challenging single-arm control paradigm with C-STEM, representing a 15.8% improvement over conventional deep learning models and a 26.3% increase above chance. Improvements in free movement metrics further corroborate these performance gains between models. Together, these findings demonstrate the enhanced decoding performance and subject adaptation enabled by the framework, highlighting the potential of foundation models to support intuitive control and more complex tasks in real-time BCI applications.

A main innovation underpinning the improved performance originates from the incorporation of online constraints in the pretraining and the architecture of the decoder. Deep learning techniques have been shown to improve EEG decoding through complex learning strategies [55], [56], [57], which have in turn enabled more complex control within BCI experimentation [17], [34]. As an extension of deep learning, foundation models strive to learn general representations of EEG from large datasets. This implies that useful task-agnostic characteristics can be inferred from EEG signals, which have indeed demonstrated strong utility in downstream offline tasks [40], [44], [53], [54]. However, online experimentation introduces additional constraints, particularly in latency for real-time feedback and limited signal pre-processing. Many current pretraining methods are not conducive to these restrictions, resulting in a lack of adaptability to online settings. These limitations represent a gap between the offline development of deep learning and foundation models in EEG and the online implementation and performance of these models in a closed-loop BCI. By not addressing these constraints, current EEG foundation models’ practicality in real-time, online BCI applications remain limited.

Our method bridges this gap between offline improvements and online application through the use of online constraints in the pretraining phase and architecture. This approach is significant for the field of EEG-BCI, as improvements in decoding accuracy for BCI tasks can be most impactful when they are able to be incorporated into online applications. While current EEG foundation models use temporal windows of 1 second [38], [45] or more [51], C-STEM looks towards smaller temporal windows of 200 ms, allowing for a more flexible configuration in downstream tasks. This design choice, coupled with a spectrogram reconstruction task in pretraining, allowed the model to learn smaller building blocks of EEG representations which still contained important spectral and temporal information (Figure 6B). We used these neural representations on out-of-distribution tasks such as motor execution (Supplementary Figure 7B) which were underrepresented in the pretraining set, and found improvements over conventional deep learning. This implies that unlabeled EEG contains learnable, widespread characteristics which can be used to improve unseen task decoding, even for online settings and applications.

Importantly, these representations create an advantage over other EEG foundation model representations in that these representations can be effectively used in online settings. Due to the design choices during pretraining, C-STEM is less impacted by decreasing temporal window sizes, while other current EEG foundation models do not handle the stringent constraints as well (Supplementary Figure 4B). While many offline tasks focus solely on trial-wise accuracy, the importance of these findings lies in the application to real-time control, which strives to provide feedback to subjects in a timely manner for applications such as robotic control. Our architecture reached stable prediction performance earlier in the trial window than conventional deep learning (Supplementary Figure 4A), This indicates that the C-STEM architecture can provide feedback faster than other methods, reducing latency and facilitating smooth and responsive control. Generally, a temporal window of one second can be acceptable for real-time feedback [14], [17] in continuous paradigms. However, paradigms which rely on single, dynamic movements may see a more compounded effect from the latency. Thus, the lower-latency characteristic, demonstrated in the C-STEM framework, is essential for BCI decoders to be practical in real-time, dynamic control applications.

Another main contribution of the study is the demonstration of the ability from the C-STEM framework to enable single-arm, dynamic movement control on a demanding task. Using C-STEM resulted in significant improvement over the conventional deep learning in both the simpler guided task (Figure 3A) and the more complex free movement task (Figure 3B). This suggests that the neural representations learned by foundation models can translate to better disentanglement of spatially and temporally complex EEG signals. Additional analysis in the per-class performance of the models (Figure 4, Supplementary Figure 2, Supplementary Figure 3) reveals that the strongest confusions were between left/right movements and up/down movements, indicating that movements within the same plane are more entangled spatially [58] and may cause more difficulty in decoding. Differences between the model confusion matrices (Figure 4B) indicate that much of C-STEM’s improved prediction accuracy arises from better disentanglement of left/right and up/down confusions, as well as moving away from the single-class default prediction that often characterizes poorly trained deep learning classifiers. This improvement suggests that C-STEM may not provide a uniform improvement across all classes, instead addressing biomechanical and model-specific issues. Overall, the improvement over the conventional deep learning methods implies that foundation model frameworks can advance the field of EEG-BCI towards two dimensions in parallel: the complexity of control mechanism and the difficulty of the online task.

Importantly, the C-STEM framework was also found to facilitate subject learning during the task. Through mixed model-data analyses, we find that data collected with C-STEM was more discriminable, allowing higher performance by EEGNet, and that this effect was reversed for data collected with EEGNet (Supplementary Figure 5B). This suggests that in addition to C-STEM’s stronger decoding ability over EEGNet, C-STEM also encourages subjects to generate a more discriminable data distribution. This implies that an online foundation model aligns better with the subject’s modulation of activity, eliciting a co-adaptive effect during the online decoding task. Further evidence comes from the performance improvement observed with mid-session finetuning (Figure 5A), where the C-STEM framework achieves stronger performance by incorporating information from the same-session distribution, although this effect does not extend to all subjects (Figure 5B). Together, these results suggest that the benefits of the foundation model framework extend beyond decoding ability, to facilitate stronger subject adaptation to the control mechanism during BCI. This finding further demonstrates the value of incorporating increasingly complex models such as C-STEM into BCI applications, as tasks become more difficult and require more subject attention.

While this study presents improvements from the C-STEM framework for more complex BCI applications, several limitations should be considered. One such limitation lies in participant selection, as all subjects were able-bodied adults with previous experience using EEG-BCI. By selecting subjects who had previously completed motor imagery EEG-BCI experiments, the sample is more likely to generate strong MI signals, potentially leading to improved performance. This behavior may not be representative of the broader able-bodied population, which likely exhibits a wider range of performance. In the general population, approximately 15-30% of individuals are unable to generate strong motor imagery signals, exhibiting the phenomenon of BCI illiteracy [59] during motor imagery tasks. Online results indicate that on average, subjects performed above chance level regardless of the algorithm used; however, it remains unclear whether this performance is attributable to the subjects’ prior EEG-BCI experience or to the complexity of the paradigm. It is also unknown whether the findings of this study would extend to clinical populations such as stroke survivors or individuals with paralysis, as these groups were not strongly represented in the pretraining dataset. Future work may therefore investigate the impact of foundation model frameworks for decoding in online experiments involving BCI-naïve subjects, BCI non-responders, or participants from clinical populations.

Another limitation lies in the design of the proposed C-STEM framework, which contains a relatively small number of parameters, compared with foundation models in other modalities, such as large language models. An EEG foundation model focused solely on motor imagery is inherently limited by the amount of available open-source motor imagery EEG data. Model components— such as the number of embeddings stored in the codebook, the dimensionality of the latent embedding space, and the number of transformer blocks in the encoder—could potentially be increased with substantial expansion of the pretraining dataset. Accordingly, one direction for future work would be to aggregate a larger corpus of EEG motor imagery data, which could increase both the diversity of the dataset and the complexity of the model, thereby further improving downstream performance. A challenge in this effort would be aligning data from the multitude sources and ensuring that important characteristics, such as sampling rate, are properly standardized across the dataset.

In this study, we used EEGNet as a representative for conventional deep learning architectures, as it has been widely evaluated in BCI research by many groups and has further demonstrated efficiency as a small, personalized model in online BCI experimentation. Future investigations may also examine deep learning architectures with increased complexity or transfer learning techniques to determine whether the current findings hold across variation in deep learning approaches.

## CONCLUSION

Studies in EEG-BCI have increasingly sought to achieve more intuitive control of external devices in complex tasks. In this study, we propose a foundation model architecture (C-STEM) pretrained on a large corpus of motor imagery data, with spectrogram reconstruction and online constraints guiding the pretraining process. We evaluate this model on demanding cursor control tasks using a single-arm, dynamic, directional control scheme. The model is evaluated in online experiments alongside a conventional deep learning architecture for comparison. Our results show that C-STEM achieves a final online accuracy of 51.3% in the four-class guided task, representing a 26.3% increase from the chance level of 25% and a 15.8% increase from the conventional deep learning EEGNet model. In addition, C-STEM demonstrates higher completion rates and lower completion times during a free-movement task. These results demonstrate the promise of EEG foundation models for online experimentation and represent a step towards enabling more complex control paradigms in motor imagery-based EEG-BCI systems.

## Author contributions

Conceptualization: M.K. and B.H. Methodology: M.K., H.W, and B.H. Data Collection: M.K., H.W., Z.J., and Y.D. Formal data analysis and visualization: M.K. Original manuscript writing: M.K. and H.W. Reviewing and editing: M.K., H.W., and B.H. Funding and resource acquisition: B.H. Supervision: B.H.

