## Supplemental Figures for "EEG Foundation Model Improves Online Directional Motor Imagery Brain-computer Interface Control"

### SUPPLEMENTARY FIGURES

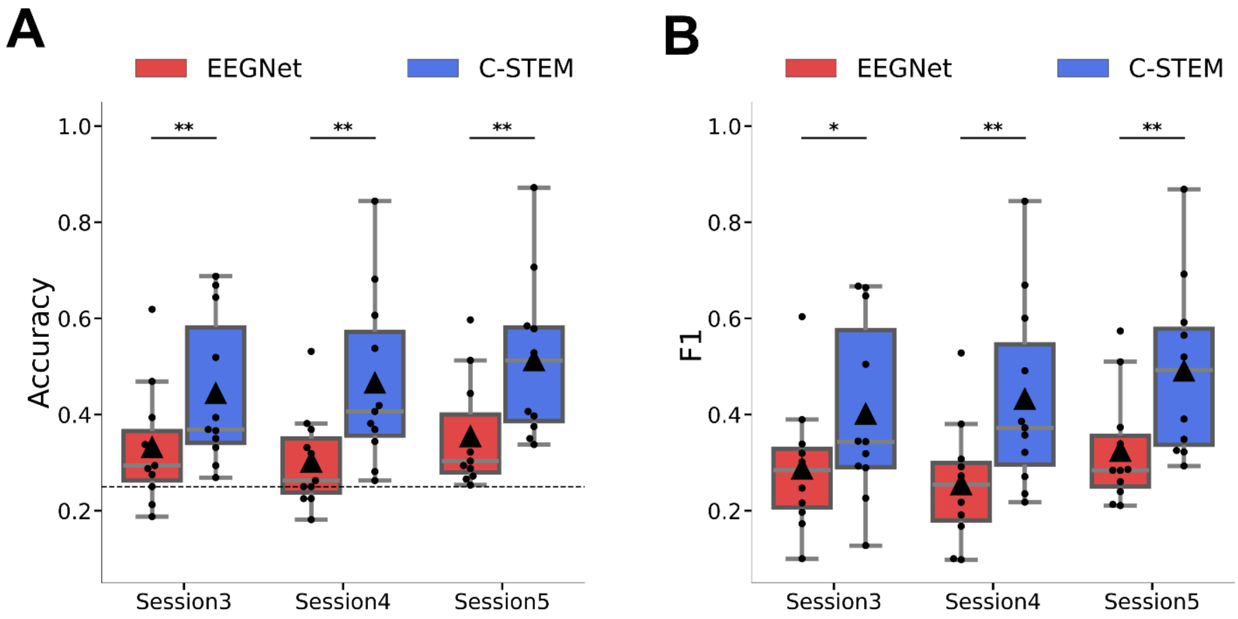


**Supplementary Figure 1.** Box-plot comparisons of Accuracy and F1 metric across models and sessions, displaying average values over subjects for the guided movement task. Center lines indicate median values, triangle markers indicate mean values, and boxes extend from the lower to upper quartile. Significance testing results between groups are displayed above bars (*** for p<0.001, ** for p<0.01, * for p<0.05, n.s. if no significance found).

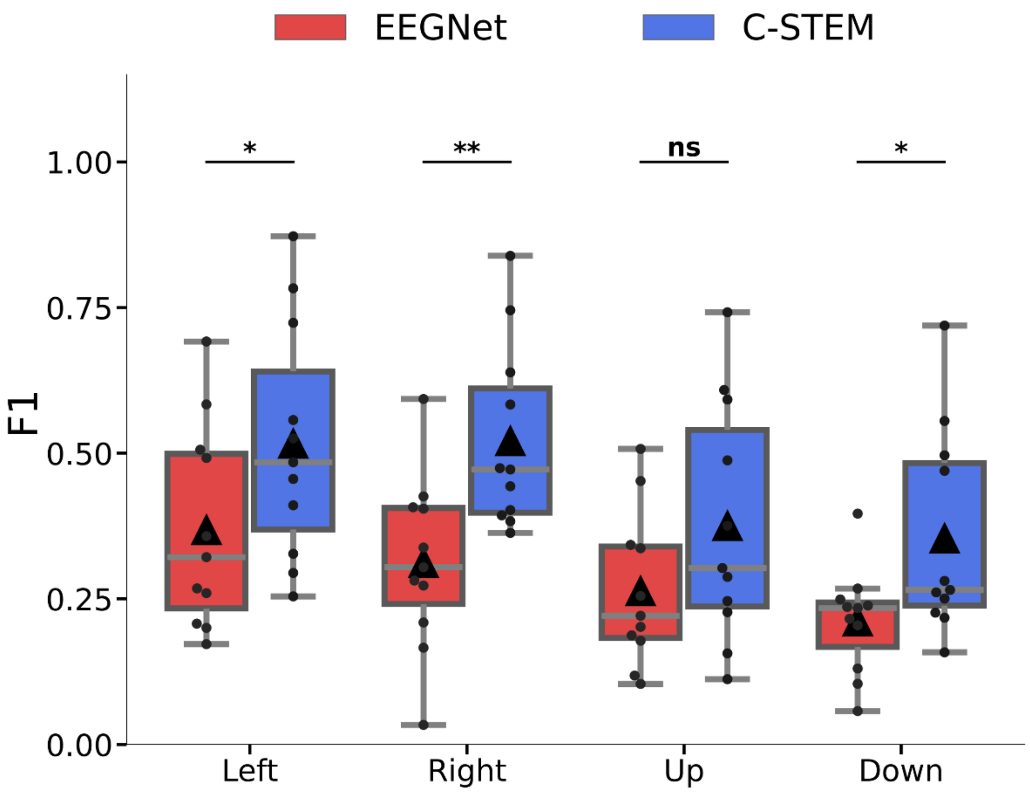


**Supplementary Figure 2.** Box-plot comparisons of the F1 metric across models displaying average values over subjects and sessions for the guided movement task divided into the four classes. Center lines indicate median values, triangle markers indicate mean values, and boxes extend from the lower to upper quartile. Significance testing results between groups are displayed above bars (*** for p<0.001, ** for p<0.01, * for p<0.05, n.s. if no significance found).

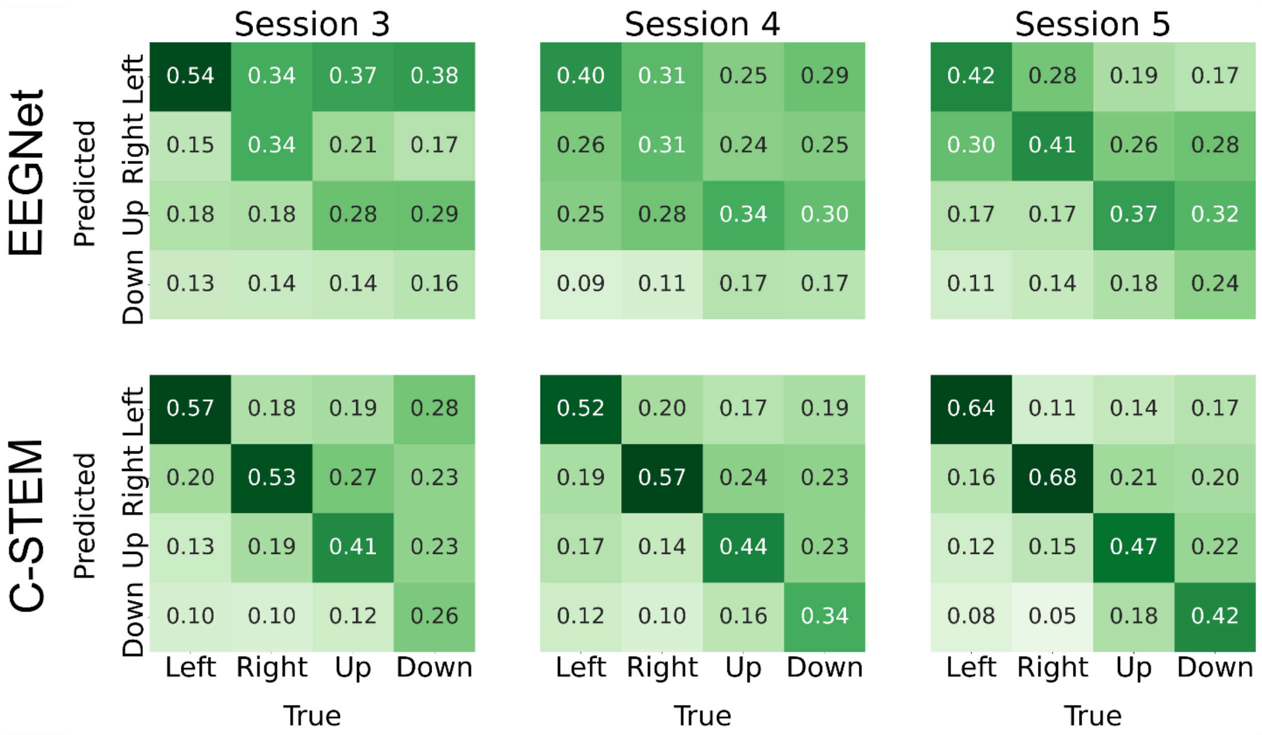


**Supplementary Figure 3.** Session-wise confusion matrices visualized for both models. The matrices provide a more granular insight into how the various models performed for each session, showing behavior and confusions that may be smoothed out by the averaging done over sessions.

**A**

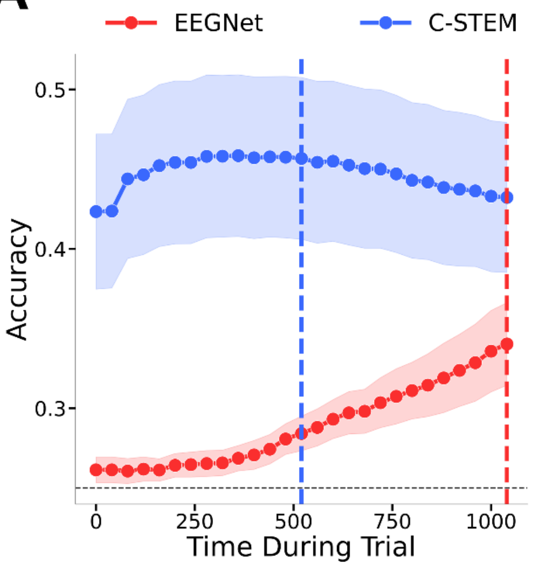


**B**

| Model | Accuracy at 200ms | Accuracy at 800ms | Accuracy at 1000ms |
|---|---|---|---|
| LaBraM | 27.6% | 28.8% | 31.0% |
| NeuroGPT | 26.5% | 32.3% | 50.8% |
| C-STEM | **36.3%** | **46.7%** | **55.8%** |

**Supplementary Figure 4.** Results of the offline explorations into latency using the C-STEM. A) Visualization of sliding windows over the duration of the entire trial. Aggregated predictions of each model are plotted over the total trial time, and peak accuracy times are marked by vertical lines. B) Comparison between alternate open-source EEG-FM models LaBraM [38] and NeuroGPT [51]. LaBraM uses a pretraining window of 1000 ms, NeuroGPT uses a pretraining window of 2000 ms, and C-STEM uses a pretraining window of 200 ms. Evaluation is done with different temporal window sizes to investigate the model performance against the constraint of temporal windows. Values are averaged over the online session data of all subjects.

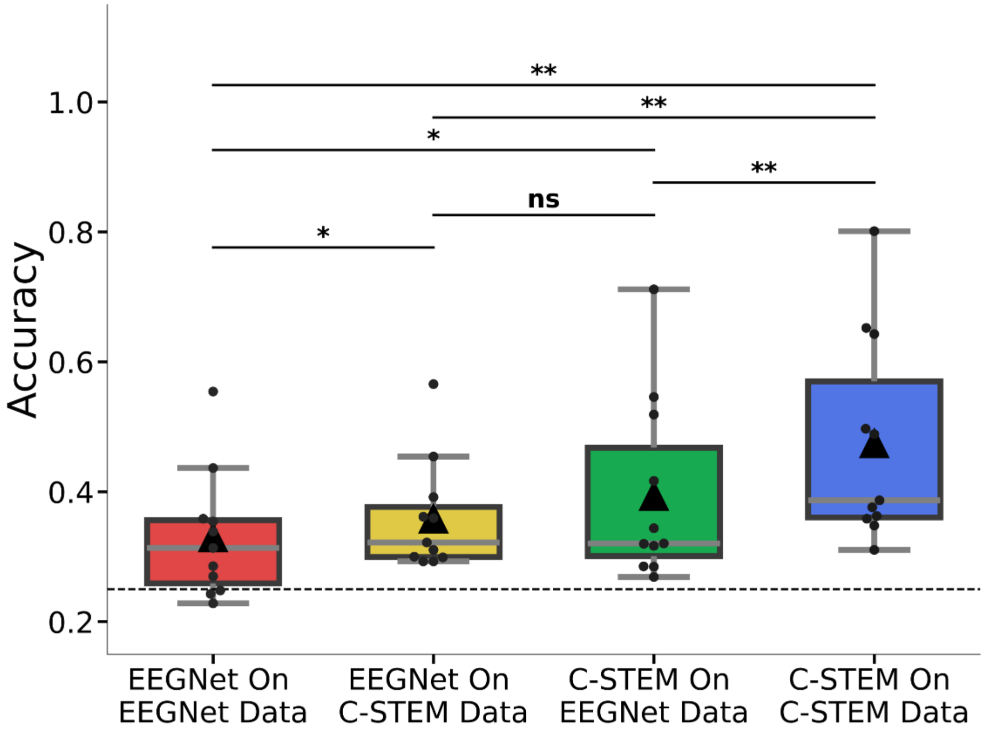


**Supplementary Figure 5.** Results for disentangling model performance and subject guidance during learning. Box-plot visualizations of the simulated accuracies of the EEGNet and FM models on runs collected with the models during the online sessions. Average values are displayed for each subject. Center lines indicate median values, triangle markers indicate mean values, and boxes extend from the lower to upper quartile. Significance testing results between groups are displayed above bars (*** for p<0.001, ** for p<0.01, * for p<0.05, n.s. if no significance found).

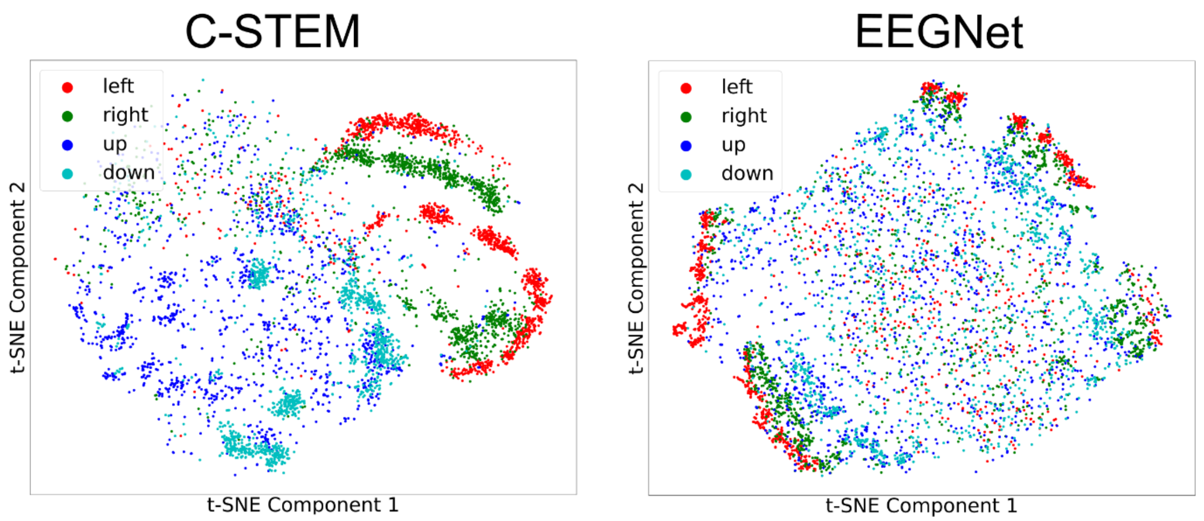


**Supplementary Figure 6**. Visualization of the embeddings generated by the proposed C-STEM and by the conventional EEGNet in downstream 4-class tasks. This comparison shows the separability of the embeddings that are used for classification.

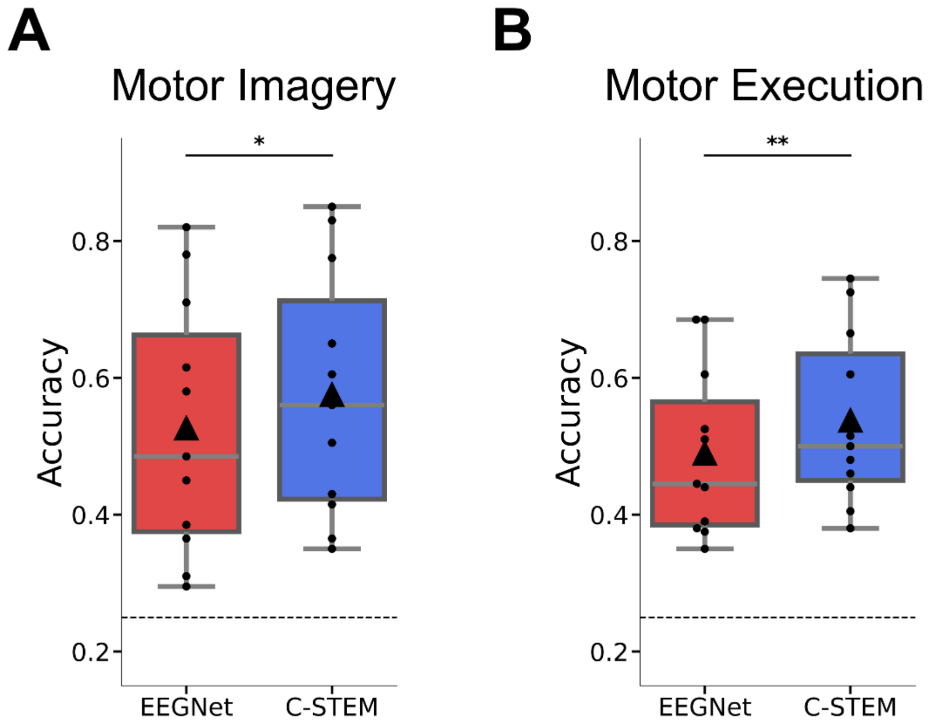


**Supplementary Figure 7.** Pseudo-online evaluation of the data in the offline sessions (Session 1,2). A) Box-plot visualizations of the pseudo-online accuracies of the EEGNet and foundation model on runs collected during motor imagery. Center lines indicate median values, triangle markers indicate mean values, and boxes extend from the lower to upper quartile. Significance testing results between groups are displayed above bars (*** for p<0.001, ** for p<0.01, * for p<0.05, n.s. if no significance found). B) Box-plot visualizations of the pseudo-online accuracies of the EEGNet and foundation model on runs during motor execution.
